# Anxiety-associated behaviors following ablation of *Miro1* from cortical excitatory neurons

**DOI:** 10.1101/2025.01.22.634105

**Authors:** Abigail K. Myers, Madison Sakheim, Cole Rivell, Catherine Fengler, Layla Jarrahy, Rachel Shin, Megan Case, Caroline Chapman, Leah Basel, Slade Springer, Nicholas Kern, Jennifer Gidicsin, Ginam Cho, Sungjin Kim, Mourad Tighiouart, Jeffrey A. Golden

## Abstract

Autism spectrum disorder, schizophrenia, and bipolar disorder are neuropsychiatric disorders that manifest early in life with a wide range of phenotypes, including repetitive behavior, agitation, and anxiety (American Psychological Association, 2013). While the etiology of these disorders is not completely understood, recent data implicate a role for mitochondrial dysfunction. To function optimally mitochondria must translocate to metabolically active intracellular compartments to support energetics and free-radical buffering; failure to achieve this localization results in cellular dysfunction (Picard et al., 2016). Mitochondrial Rho-GTPase 1 (*Miro1*) resides on the outer mitochondrial membrane and participates in neuronal microtubule-mediated mitochondrial motility and homeostasis (Fransson et al., 2003). Previous research implicates the loss of *MIRO1* as a contributor to the onset/progression of neurodegenerative diseases including amyotrophic lateral sclerosis, Alzheimer’s disease, and Parkinson’s disease (Kay et al., 2018). We have hypothesized that MIRO1 also has a role in nervous system development and function (Lin-Hendel et al., 2016). To test this, we ablated *Miro1* from cortical excitatory progenitors by crossing floxed *Miro1* mice with *Emx1-cre* mice. We found that mitochondrial mis-localization in migrating excitatory neurons was associated with reduced brain weight, decreased cortical volume, and subtle disruptions in cortical organization. Adult *Miro1* conditional mutants exhibit agitative-like behaviors, including decreased nesting behavior and abnormal home cage activity. Open field testing revealed anxiety-like behavior and elevated plus maze and wide/narrow box testing found the mice avoided confined spaces. Our data link MIRO1 function with mitochondrial dynamics in the pathogenesis of several neuropsychiatric disorders and implicate mitochondrial localization in anxiety-like behaviors.

**Significance:** Neuropsychological disorders such as autism spectrum disorder, schizophrenia, and bipolar disorder have overlapping symptoms and behaviors. While the mechanisms underlying these disorders are not completely understood, recent evidence suggests mitochondrial dysfunction and mis-localization within a cell could play a role. Mitochondria are organelles that provide energy and other self-regulating processes to the cell. Previous research from our lab has shown distinct dynamic localization patterns within migrating excitatory and inhibitory neurons may be important during development. To further examine the importance of mitochondrial localization, we ablated MIRO1, a protein important for coupling mitochondria to motor proteins, in excitatory neurons. Mitochondria mis-localize in migrating excitatory neurons, and this is associated with a loss of motor skills and anxiety-like behavior in post-natal mice.

## Introduction

Neuropsychiatric disorders comprise a heterogenous group of complex conditions affecting as many as 1 in 5 individuals in the U.S (National Institute of Mental Health, 2023). Their etiologies remain heterogenous and poorly understood, though many appear to originate prenatally (Chan, 2006). These disorders can manifest early in life with a wide range of behavioral phenotypes that may include repetitive behavior, agitation, resistance to touch, psychosis, and anxiety, along with a variety of other features (Miyoshi et al., 2010).

While the etiology of some neuropsychiatric disorders is believed to be multifactorial, recent data suggest mitochondrial dysfunction could play a role in both syndromic and non-syndromic cases (Pei and Wallace, 2018). Population-based studies of individuals with a neuropsychiatric disorder indicate a prevalence of mitochondrial dysfunction as high as 80% in disorders such as autism spectrum disorder, implicating a role for mitochondria during brain development (Oliveira et al., 2005; Giulivi et al., 2010; Siddiqui et al., 2022). Mitochondria play a role in neuronal migration, circuit formation, synaptogenesis and plasticity and given that these processes have been implicated in the development of some neuropsychiatric disorders, we have postulated that mitochondrial dysfunction would affect developmental processes resulting in post-natal behavior phenotypes (Chan, 2006; Rahman, 2012; Lin-Hendel et al., 2016; Fame and Lehtinen, 2021).

Mitochondria are double – membrane - bound organelles essential for normal cellular physiology. They produce ATP, buffer ion homeostasis, assist with free-radical elimination, support autophagy, and assist in lipid metabolism (Nunnari and Suomalainen, 2012; Mahadevan et al., 2021). To fulfill these various roles, mitochondria must shuttle to different intracellular compartments. Mitochondrial Rho-GTPase *(Miro1)*, is a motor adaptor protein that links mitochondria to molecular motors for transport along microtubules (Tang, 2015; López-Doménech et al., 2018). In addition to transportation, *Miro1* is known to help coordinate mitochondrial movement, fission, fusion, and mitophagy (Hsieh et al., 2016). In mature neurons, mitochondrial movement is essential for maintenance of dendritic branches, synapse formation and maintenance, and proper synaptic transmission (López-Doménech et al., 2016). The loss of *Miro1* has been associated with neurodegenerative diseases such as Parkinson’s disease, Alzheimer’s disease, and amyotrophic lateral sclerosis (Zhang et al., 2015; Grossmann et al., 2019, 2020). MIRO1 also interacts with PINK1, Parkin, α-synuclein, and LRRK2 to mediate mitophagy, preserving mitochondrial quality in neurons (Lin and Sheng, 2015; Shaltouki et al., 2018; Grossmann et al., 2020). Dysregulation of these protein interactions has been implicated in cerebral cortical, brainstem, and hippocampal neuronal degeneration resulting in spasticity, weakness, and memory loss (Nguyen et al., 2014).

In addition to its role in mature neurons and neurodegeneration, recent data implicate *Miro1* in the onset of several neuropsychiatric diseases. For example, *Miro1*-deficient cortical neurons fail to develop normal dendritic arbors (López-Doménech et al., 2016) and MIRO1 seems to form a complex with Disrupted-in-Schizophrenia-1 (DISC1), a protein that is important for neurite outgrowth and has been associated with a wide array of neuropsychiatric disorders (Ogawa et al., 2014; Norkett et al., 2017). A sequence variant of DISC1, R37W, was associated with individuals diagnosed with schizophrenia, depression, and anxiety in several families and was found to perturb anterograde mitochondrial transport in neurons (Ogawa et al., 2014).

Given the relationship between neuropsychiatric disorders and cerebral cortical neuronal dysfunction (McManus and Golden, 2005; Rubenstein, 2011), we hypothesized that disruption of *Miro1* in developing excitatory neurons would result in behavioral disorders. The conditional abrogation of *Miro1* from projection neurons resulted in agitative-like behaviors including resistance to touch, decreased nesting behavior, repetitive behaviors, and decreased interactions with littermates. The mice also have abnormal behavior during open field testing and other anxiety-like behaviors. These data suggest *Miro1* is required for the normal development of excitatory neurons and provide novel insights into the underlying pathogenesis of neuropsychiatric diseases that include anxiety-like behaviors.

## Materials and Methods

### Mice

Mice were housed and sustained in the Animal Vivarium at Hamilton College (initial studies were conducted at the Brigham and Women’s Hospital) and were given food and water ad libitum. All experiments were approved by the Institutional Care and Use Committee at Hamilton College and Brigham and Women’s Hospital. *Miro1(^f/f^)* mice (Strain #:031126, The Jackson Laboratory) and *Emx1-cre(^+/+^)* mice (Strain #005628, The Jackson Laboratory) were maintained on a C57/Bl6 background. *Miro1(^+/+^)*, *Miro1(^f/+^)*, *Miro1(^f/f^)*, *Miro1(^+/+^); Emx1-cre(^+/+^)*, and *Miro1(^+/+^); Emx1-cre(^+/-^)* mice were littermates of the *Miro1* conditional mutant mice and used as controls for experiments. *Miro1* conditional mutant mice were generated across six different litters.

### Weight Measurements

Mice were weaned on postnatal day 21 (P21) and fed a lab diet (5058, LabDiet). All mice were genotyped and weighed every 7 days following weaning for 5 weeks. The percentage difference in weight between controls and knockout animals was calculated by dividing the average weight of the *Miro1* conditional knockout animals by the average weight of the control animals during that week. Adult mice (control and knockout) were weighed between 6 to 10 months of age to examine weight differences in adult animals.

### Food Intake

During cage changes, food was weighed out for individual mice for one week and added to the food trough. At the end of the week prior to the next cage change, the remaining food in the trough was weighed to establish food intake.

### Daily cage recordings

Control and *Miro1* conditional mutant mice were singly housed in clear (18.5 cm x 21 cm) cages. Red light was used to visualize the mice without disrupting their light/dark cycles. Cameras (WiFi Indoor Camera G7, Galayou Smart Home Security Cameras) were positioned on the top of each cage to monitor the movements of the mouse. Each cage was recorded during the mouse dark cycle when they are most active. ANY-MAZE software (Stoelting Co.) was used to track each pre-recorded movie for two hours between 12 am and 2 am during the dark cycle.

### Hindlimb Footprint Pattern Test

Receipt paper was placed on and secured to the floor in a narrow hallway (110 cm x 10 cm). The hindlimb feet of adult mice were brushed with India Ink (Speedball) using a paintbrush. The mice were placed on the receipt paper in the narrow hallway and allowed to walk one length of the hallway. A series of three consecutive footprints for each hindlimb were measured.

### Forelimb Grip Strength Test

A T-shaped pull bar was attached to a force meter (Ametek Chatillon DFS II, 10 N) fixed to the table. Mice were held by the tail and allowed to grasp the pull bar with their forelimbs. Once the mouse had a good grip on the pull bar the mouse was gently pulled by the tail until they let go of the bar. The force was recorded (in Newtons) on three consecutive trials with each mouse.

### Vertical Pole Test

A wooden cylindrical beam of 92.2 cm in length and 1.8 cm in diameter was used to assess gross motor control of body muscles utilized in hanging onto the thick beam as opposed to grip strength alone. The wooden beam was fixed at one end in the middle of a large 1.02 x 1.02-meter rodent testing container. The mouse was placed upon the wooden beam which was then slowly raised from one end until an angle of 90 degrees was reached. Upon reaching 90 degrees, the time to fall off the pole was measured (maximum 60 seconds). Mice that were unable to stay on the beam during the elevation phase were given a 2-minute rest and then retested. If a mouse was unable to reach 90 degrees elevation during the retest, the final degree of elevation was recorded.

### Hanging Wire Test

Mice were placed on top of a 26 cm by 36 cm wire grid that had 1.2 cm^2^ square holes. The wire grid was lightly shaken horizontally for the mice to grasp the wire with all paws, then flipped 180 degrees and held 20.0 cm high from a testing area. The mice were then timed while inverted with a maximum threshold of 60 seconds.

### Wire Grid Test

The wire grid was placed on top of a box and a camera placed on the floor of the box to record the limb and paw placement throughout the test. The mouse was placed on the wire grid and then a clear plastic container was used to restrict the mouse to the wire grid area. Each mouse was tested for five minutes on the wire grid. The video was analyzed by counting the number of times a paw lost grip and fell through the wire grid.

### Tests for anxiety-like behaviors

Before each anxiety-like behavior test, cages were transported from the animal facility to the behavior room and allowed to acclimate to the new environment for 15 minutes. Tests were completed under normal room lighting and with general room background noise. At the conclusion of each trial the apparatus was cleaned with a 70% ethanol solution.

### Open Field Test

Adult mice were placed in the center of a (30 cm x 40 cm) arena and allowed to travel freely for 5 minutes. The arena was divided into a central region and an outer region to distinguish locations within the box. Movements around the box were tracked using the ANY-MAZE software package (Stoelting Co.). Distance, speed, time mobile, freezing episodes, center entries, and distance traveled in the center were exported as measurements from the ANY-MAZE software. A freezing episode was defined as no movement except for respiration for a period of 250 ms.

### Elevated Plus Maze

Adult mice were placed in the center of the elevated plus maze at the start of the trial and allowed to explore the maze for 5 minutes. The ANY-MAZE software package (Stoelting Co.) was used to label each wing of the elevated plus maze as an “open” or “closed”. The time and distance traveled in the open and closed arms was saved from the software.

### Wide/Narrow Box Test

Developed as a measure of ‘claustrophobia’, a 60 cm x 60 cm box was constructed and divided into two 30 cm sections: a wide section (60 cm width x 30 cm length) and a narrow section (5 cm width x 30 cm length; adopted from El-Kordi et al., 2013). The wide section of the box was lit to 300 lumens (lx) and the narrow section of the box lit to 150 lx. Mice were placed in the wide section of the box facing the narrow section to begin the trial and were allowed to roam freely in the box for 10 minutes. The ANY-MAZE software (Stoelting Co.) was used to track the path of the mice in the box and measure the time spent in the narrow and wide sections of the box.

### Plasmid Construction

Separate fragments of EGFP, mito-DsRed2 (TaKaRa, Cat #:632421) and P2A derived from porcine teschovirus-1 2A (Kim et al, 2011) were amplified by PCR. The fragments were subcloned into pCAG (Addgene) by Geneart (Thermofisher). The mito-targeting sequence “SVLTPLLLRGLTGSARRLPVPRAKIHSL” was derived from the precursor of subunit VIII of human cytochrome C oxidase (Rizzuto et al., 1989).

### Electroporation and slice culture experiments

Mouse embryos were harvested on embryonic day (E)14.5. Brains were immediately removed and placed into cold complete HBSS bath (Tucker et al., 2006; Lysko et al., 2014). The *pCAG-GFP-P2A-mitoDSred* plasmid was micropipetted into both lateral ventricles and electroporated into the ventricular zone of the cerebral cortex (Nepa Gene CUY21 electroporator; 45V, pulse interval: 100ms, pulse duration: 100ms, number of pulses: 4). Brains were embedded in 3% low melting point agarose (Fisher Scientific, Cat # BP165-25) and cut into 300μm sections (Leica VT 1000S). Slices were placed on transwell inserts (BD Biosciences, Cat #:353102) coated with laminin/poly-L-lysine and cultured for 3 days (Polleux and Ghosh, 2002). Slices were fixed on the third day with 4% paraformaldehyde.

### Immunohistochemistry

Adult mice were anesthetized and perfused with 1x PBS followed by 4% paraformaldehyde. Brains were harvested, post-fixed in 4% paraformaldehyde for 24 hours, and then placed in 30% sucrose to prepare for cryosectioning. Each brain was sectioned in 60 μm intervals on a freezing microtome (Leica SM 2000R). Embryonic brains were harvested at E13.5 or E15.5, and then placed into 10% sucrose overnight, followed by 30% sucrose the next day for 24 hours to prepare for cryosectioning. Brains were sectioned at 15μm on a cryostat (Leica CM1950 or Dakewe 6250). Sections were blocked with 5% normal goat serum. Adult tissue was labeled with Rabbit anti-CUX1 (1:250, Proteintech Cat #: 11733-1-AP) and Rat anti-CTIP2 (1:250, Abcam, Cat #: ab18465), and embryonic tissue was labeled with Rat anti-KI67 (1:500, ThermoFisher, Cat #: 14-5698-82), followed by Alexa Fluor secondary antibodies (Invitrogen, Goat anti-Rabbit Alexa Fluor 488 Cat #: A-11008 and Goat anti-Rat Alexa Fluor 555 Cat #: A-21434) with Dapi labeling, and imaged on a Leica SP5 or Zeiss 910 Confocal Microscopes or Leica DM6 B Microscope.

### Analysis of mitochondria location within cell body

The somas of migrating excitatory neurons were segmented and vectors were calculated with the Aivia software (Leica Microsystems). A spherical coordinate system with cartesian coordinates was used to measure where the mitochondria labeled with MitoDSred were localized within the cell body.

The following scripts in the measurement component of the Aivia software were used to measure the angular location of the mitochondria within the soma using the azimuth, the angle within the x-y plane: *X Displacement*: ValueAtFrame([Centroid X (Mitochondria)], 2) - ValueAtFrame([Centroid X (Mitochondria)], 1), *Y Displacement:* ValueAtFrame([Centroid Y (Mitochondria)], 2) - ValueAtFrame([Centroid Y (Mitochondria)], 1), *Z Displacement:* MaxOverTime([Centroid Z (Mitochondria)]) - MinOverTime([Centroid Z (Mitochondria)]), *Azimuth:* Atan2([Y Displacement (Mitochondria)], [X Displacement (Mitochondria)]) * (180/PI()).

### Statistical analysis

A Student’s t-test was used to compare control and *Miro1^CKO^*groups in our brain weight and behavior analyses. Two-way analysis of variance (ANOVA) testing was used to analyze the bin distributions of cells in the cortex. The Watson-Wheeler Test was used to compare circular data for mitochondrial localization within the cell body.

The longitudinal data of behavior components from the elevated plus maze and open field tests were analyzed using a rank-based nonparametric method (Noguchi, K., Gel, Y. R., Brunner, E., Konietschke, 2012) to examine treatment group (*Miro1^CKO^*vs. control), time (10 to 300 seconds), and their interaction effects on individual behavior outcomes. For speed data from the elevated plus maze, missing data were imputed by the mean value of non-missing data at given timepoint in each group. Analyses were performed using R version 4.2.3 (R Core Team, 2023; circular and nparLD packages) with two-sided tests at a significance level of 0.05.

## Results

### Miro1^CKO^ mice have a reduced body and brain size

*Miro1^f/f^* mice were crossed with *Emx1-cre^+/+^*to generate *Miro1^+/-^;Emx1-cre^+/-^* that were then crossed to generated *Miro1^-/-^;Emx1-cre^+/-^* (subsequently referred to as *Miro1^CKO^* or CKO) and littermates with the genotypes listed in the Materials and Methods. By weaning, *Miro1^CKO^* mice were noticeably smaller than littermate control mice, weighing 22.6% less (Figure 1A). By 8 postnatal weeks, the difference was less pronounced (11.9%) with a 5% reduction in average weight for adult *Miro1^CKO^* mice. To determine if food intake accounted for the weight reduction the average daily weight of food consumed was measured. Over seven days there was no statistical difference in food consumptions (31.2 vs 32.8 g for control and *Miro1^CKO^* mice respectively). *Miro1^CKO^*mice were also found to live a full lifespan of 1.5 years or longer. Furthermore, the *Miro1^CKO^*brains weighed less than littermate control brains (Figure 1C, D), and the cortical volume was reduced in adult *Miro1^CKO^* brains (Figure 1E). KI67 labeling at E13.5 revealed a statistically significant shift in the percentage of cycling cells from Bin 3 (lower cortical plate) to Bin 8 (ventricular zone; Figure 1F, G), while there were no changes in KI67 labeling at E15.5 (Figure 1H, I).

**Figure 1:**
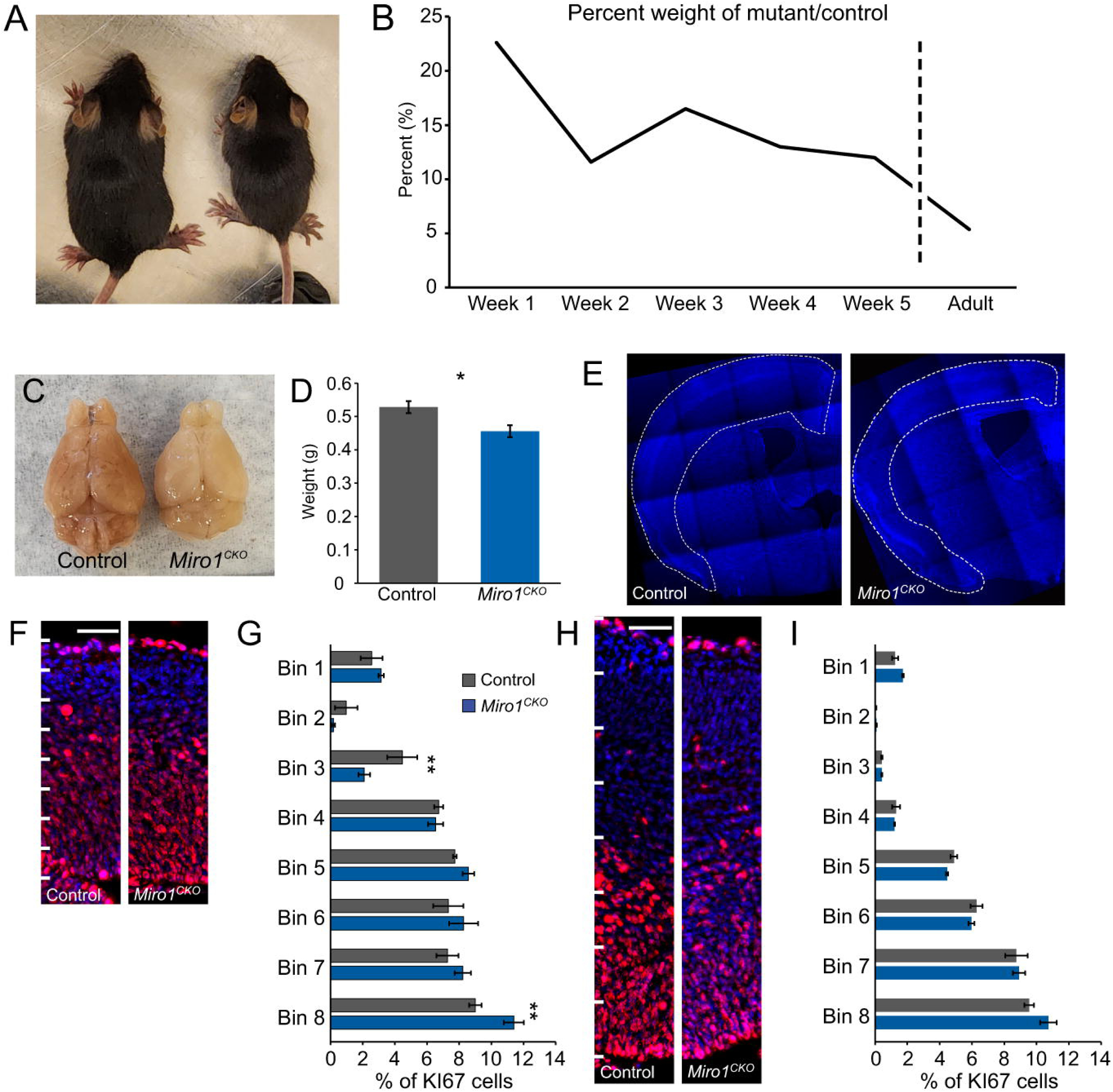
*Miro1^CKO^*body and brain size are small. **A,** *Miro1^CKO^* mice (right) are smaller than their littermate controls (left). **B,** Quantification of difference in weight between controls (weaning age: n=9, adult: n=6) and *Miro1^CKO^* mice (weaning age: n=12, adult: n=8). **C,** Size differences between control and *Miro1^CKO^* brains. **D,** Quantification of brain weight (n=5, t(8)=2.824, p=0.0224, Student’s Unpaired T-Test). **E,** Coronal sections of control and *Miro1^CKO^*brains (Dapi). **F,** KI67 labeling at E13.5 (Scale Bar: 50 μm; Bin 1 – Pial Surface; Bin 8 -Ventricular Surface). **G,** Quantification of KI67 labeling in E13.5 cortices (n=3, F(7, 32) = 3.000, p=0.0154, Two-way ANOVA; Fisher LSD Post-Hoc: Bin 3, p=0.0070, Bin 8, p =0.0065). **H,** KI67 labeling at E15.5 (Scale Bar: 50 μm; Bin 1 – Pial Surface; Bin 8 -Ventricular Surface). **I,** Quantification of KI67 labeled cells in E15.5 cortices (n=3, F(7, 32) = 1.615, p= 0.1669, Two-way ANOVA). * p≤0.05, **p≤ 0.01

### Mitochondria and projection neurons are mis-localized in migrating Miro1 ^CKO^

To establish intracellular mitochondrial distribution, *pCAG-EGFP-P2A-mitoDSred* was electroporated into control and *Miro1^CKO^* mouse forebrains on E14.5, the brains were harvested and slice cultures established from them. After three days in culture the mitochondria localization was assayed in migrating excitatory neurons. Control neurons displayed their expected mitochondria localization, proximal to the nucleus in the direction of the leading process as we previously described (Figure 2A, C-Left graph; Lin-Hendel et al., 2016). In contrast, the mitochondria in *Miro1^CKO^* neurons were predominately mis-localized to the rear of the migrating cells (Figure 2B, C-Right graph).

**Figure 2:**
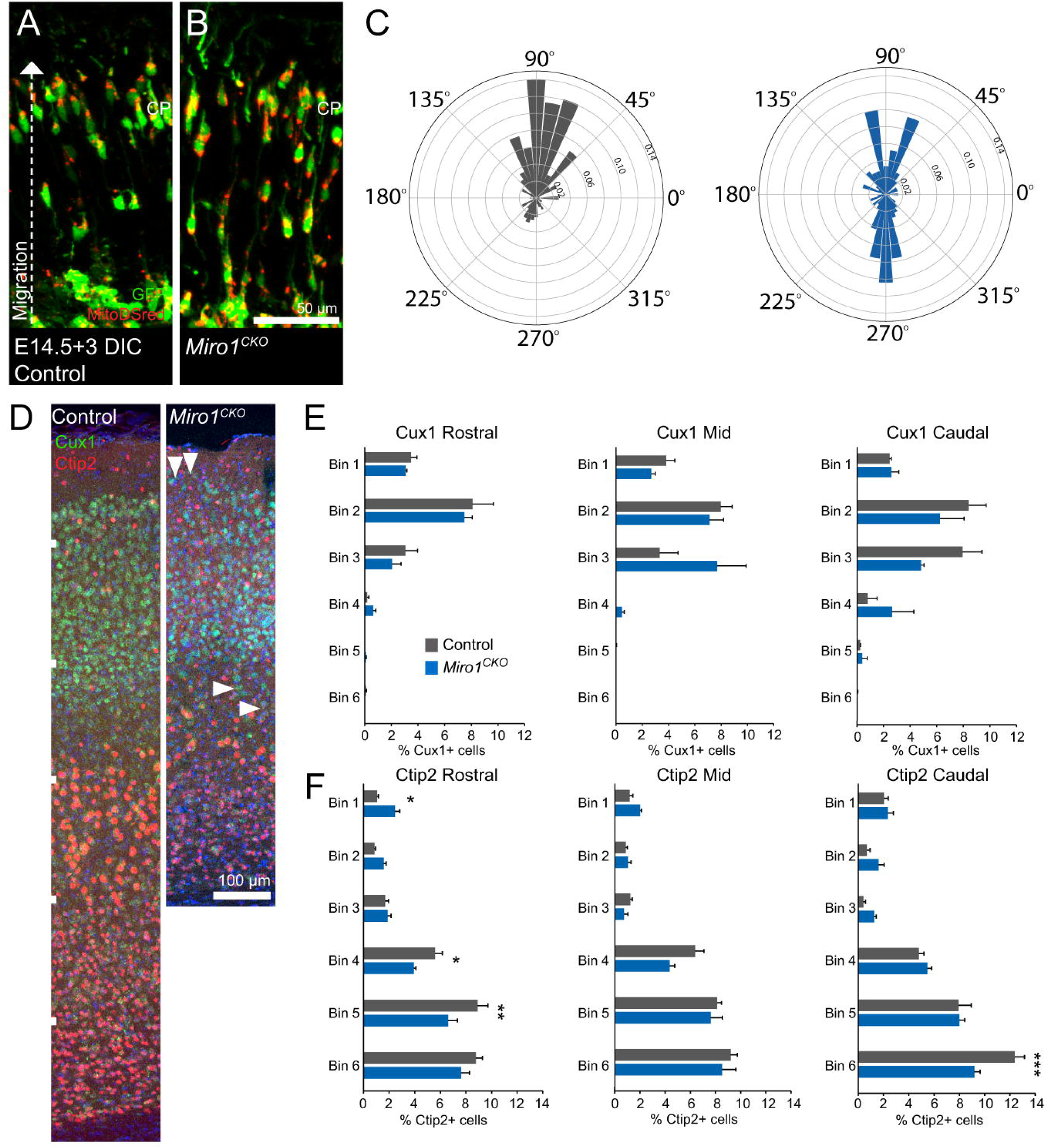
*Emx1-cre* mediated ablation of *Miro1* leads to mis-localization of mitochondria in migrating excitatory neurons and cellular changes in adult cortices. **A, B,** Migrating excitatory neurons (GFP) and labeled mitochondria (MitoDSred) in E14.5 control and *Miro1^CKO^*cortices. **C,** Quantification of red intensity (MitoDSred) location in the cell bodies of migrating excitatory neurons (Control n=265 cells, *Miro1^CKO^*n=211 cells; P<0.001, Watson-Wheeler Test for Significance). **D,** Distribution of cells in layers 2/3 (Cux1) and layers 5/6 (Ctip2) of mid coronal sections. **E,** Quantification of Cux1-labeled neurons in rostral, mid, and caudal locations (n=3, rostral: F(5, 24)= 0.3726, p =0.8624, mid: F(5, 24)= 2.581, p=0.0527, and caudal: F(5, 24)=1.782, p=0.1546; Two-Way ANOVA). **F,** Quantification of Ctip2-labeled neurons in rostral, mid, and caudal locations (n=3, rostral: F(5,24)=4.994, p=0.0028, mid: F(5, 24) = 1.894, p=0.1329, caudal: F(5, 24) =5.109, p=0.0025; Two-Way ANOVA). Post-Hoc Fisher’s LSD: n.s. P>0.05, * P≤0.05, **P≤0.01, *** P≤0.001

To determine if cortical neurons were properly positioned in the cerebral cortex, rostral, mid, and caudal coronal sections were immunolabeled with antibodies to CUX1 (labeling layers 2/3) and CTIP2 (labeling layers 5/6). Images of the cortex (Figure 2D) were divided into 6 equal bins from the pial surface to the white matter. Although the overall distribution of the cells was not overtly disrupted, minor distribution changes were noted (Figure 2D). In the rostral, mid, and caudal locations, no significant changes to the distribution of CUX1-labeled cells were observed, although in the mid-section it trended close to significance (Figure 2E). However, CUX1-labeled cells were found in regions outside of layers 2/3 in many cropped images from *Miro1^CKO^*cortices including layer 1 (Figure 2D, down pointing arrows) and layer 4 (Figure 2D, side pointing arrows). The number of CTIP2 positive cells was significantly reduced in the rostral and caudal axis of *Miro1^CKO^* cortices (Figure 2F; see rostral: Bins 1, 4, and 5 and caudal: Bin 6). Although similar trends were observed at the mid location, this did not reach significance. In each location an increase in cells can be seen in layer 1 in *Miro1^CKO^* cortices, but this only reached significance in the rostral location (Figure 2D, F).

### Abnormal home cage behaviors

*Miro1^CKO^ mice* were observed to be hyperactive and to engage in repetitive activities, including circling the periphery of the cage (Supplemental Movie 1). They also displayed poor nesting (Figure 3D-E) and resistance to being handled by animal care and research staff (biting, squealing, and extreme writhing). To determine if the hyperactive behaviors were only in response to workers being present or if they were ever-present, control and *Miro1^CKO^* mice were tracked during their dark cycle (see methods). Traces from the overnight recordings showed control mice traveled throughout their cage and spent time in their nest (Figure 3A-Left), whereas *Miro1^CKO^*mice traveled in a circular track around the cage perimeter throughout the 2-hour window, never returning to a nest (Figure 3A-Right). The distance and rate of travel found for *Miro1^CKO^* mice were significantly increased when compared to controls (Figure 3B,C).

**Figure 3:**
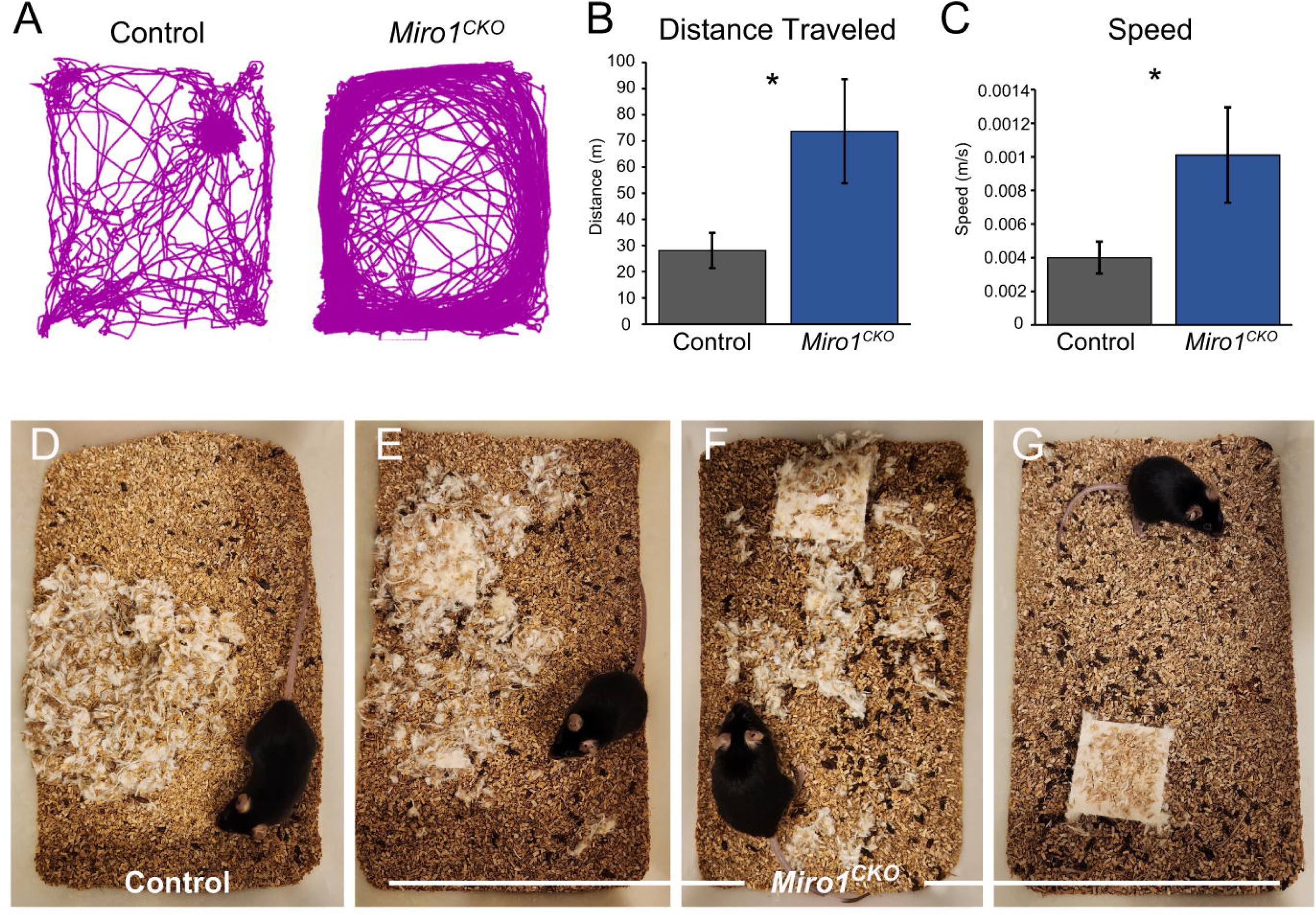
*Miro1^CKO^*mice display abnormal home cage behavior. **A,** Example traces from home cage tracking. **B,** Quantification of distance traveled in home cage (Controls n= 12, *Miro1^CKO^* n=9) **C,** Quantification of speed in home cage (Control n= 12 and *Miro1^CKO^* n= 9, Distance: t(19)=2.423, p=0.0256, Student’s Unpaired T-Test, Speed: t(19)=2.282, p=0.0342, Student’s Unpaired T-Test). **D-F,** Spectrum of *Miro1^CKO^* nesting in home cage. *P≤0.05

### Impaired Motor skills

Previous data suggested that *Miro1^CKO^* mice ablated with *Eno2-cre* develop motor neuron disease (amyotrophic lateral sclerosis)-like phenotypes (Nguyen et al., 2014). Although our mice moved freely throughout their cage without noticeable impairment, to establish if they had any subtle motor impairments, gross motor skills were assessed using the hind-limb footprint pattern test (Figure 4A). Hind-base width and stride length measurements showed no difference between the control and *Miro1^CKO^* mice (Figure 4B,C). The vertical pole test, hanging wire, wire grid, and fore-limb grip strength were used to further investigate motor coordination and strength in the *Miro1^CKO^*mice. Although most of the control and *Miro1^CKO^* mice were able to complete the initial vertical pole incline (with the exception of one control and one *Miro1^CKO^* falling off at 63° and 37° respectively), the *Miro1^CKO^* mice were unable to stay on the rod as long as the controls at 90° (Figure 4F). Additionally, the *Miro1^CKO^* mice showed statistically significant decreases in time on the hanging wire, reductions in fore-limb grip strength, and increases in foot slips on the wire grid (Figure 4D, E, G). Together these data suggest *Miro1^CKO^* mice have subtle motor skill deficits, although less severe than the *Eno2-cre*;*Miro1* conditional mutant mice.

**Figure 4:**
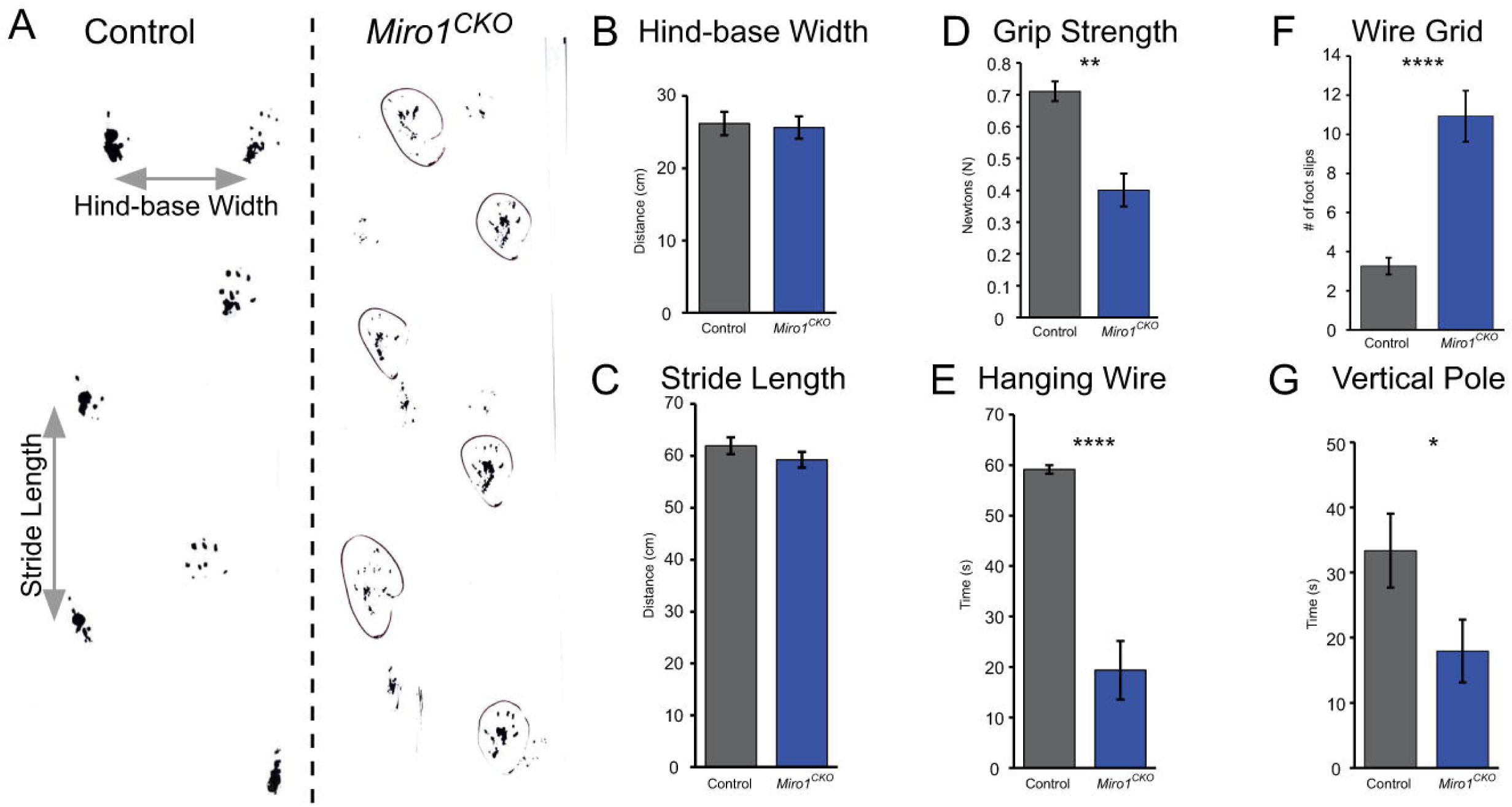
*Miro1^CKO^*mice have impaired motor strength and coordination. **A,** Examples of control and *Miro1^CKO^* footprint patterns and measurements. **B,C,** Quantification of hind-base width and stride length (Control n= 13 and *Miro1^CKO^*n= 12 Hind-Base Width: t(23)=0.2952, p=0.7705, Student’s Unpaired T-Test, Stride Length: t(23)=1.198, p=0.2433, Student’s Unpaired T-Test) from the footprint pattern test. **D-G,** Quantification of grip strength, vertical pole, and wire grid data (Grip Strength: Control n= 25 and *Miro1^CKO^*n= 11, t(34)=3.420, p=0.0016, Student’s Unpaired T-Test; Hanging Wire: Control n= 15 and *Miro1^CKO^* n= 15, t(28)=6.821, p<0.0001, Student’s Unpaired T-Test; Vertical Pole: Control n= 15 and *Miro1^CKO^* n= 15, t(28)=2.071, p=0.0477, Student’s Unpaired T-Test; Wire Grid: Control n= 15 and *Miro1^CKO^*n= 15, t(28)=5.570, p<0.0001, Student’s Unpaired T-Test). * P≤0.05, **P≤0.01, **** P≤0.0001

### Anxiety-like behavior in Miro1^CKO^ mice

To determine if the hyperactive and agitative behavior was due to anxiety, we tested the mice using two anxiety-like behavior paradigms: the open field test and the elevated plus maze. We examined components of behavior over time and by averaging. In the open field test, the average time mobile for the *Miro1^CKO^*mice was significantly lower than the controls (Figure 5B, Right graph). During the experiment, the control group had a longer time mobile compared to CKO group (p<0.001). The time mobile for the control group decreased over time (p=0.001) while there was no change in the time mobile for the CKO group (Figure 5B, Left graph; p=0.674).

**Figure 5:**
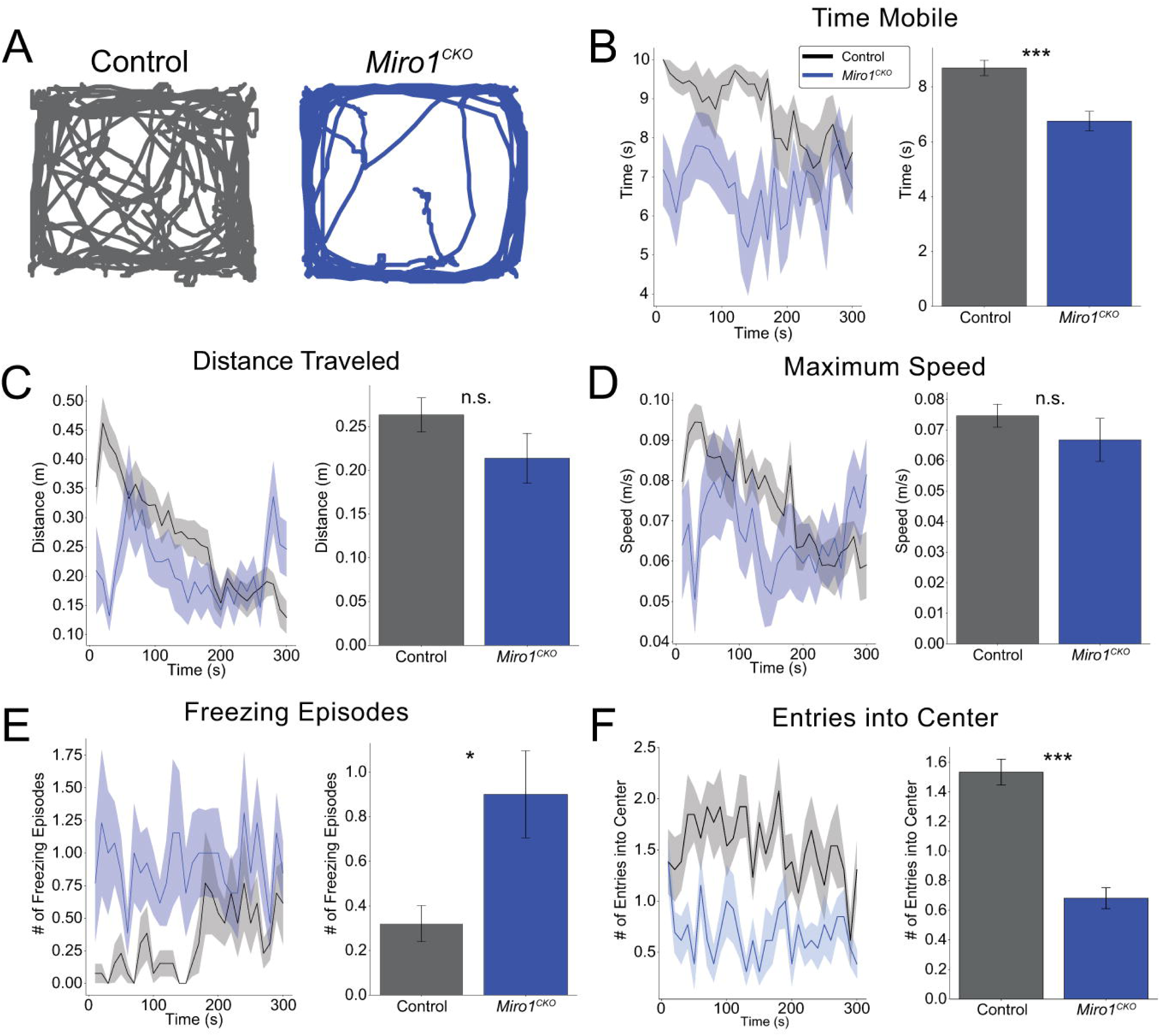
*Miro1^CKO^*exhibit anxiety-like behavior during the open field test. **A,** Example traces from the controls and *Miro1^CKO^* mice during 5-minute open field test. **B-F,** Quantification of time mobile, distance traveled, maximum speed, freezing episodes and entries into the center (Controls n = 14, *Miro1^CKO^* n = 13; Time mobile: Time mobile: t(25)=4.12, p=0.0004, Student’s Unpaired T-Test, Distance: t(25)=1.37, p=0.183, Student’s Unpaired T-Test, Max speed: t(25)=0.955, p=0.349, Student’s Unpaired T-Test, Freezing episodes: t(25)=-2.63, p=0.0145, Student’s Unpaired T-Test, Entries into center: t(25)=7.25, p=<0.0001, Student’s Unpaired T-Test). n.s. P>0.05, * P≤0.05, **P≤0.01, *** P≤0.001

The average maximum speed and total distance traveled were not significantly different between the control and the *Miro1^CKO^* mice (Figure 5C,D, Right graphs); however, the time traveling was significant (Figure 5B right graph, p<0.001). The distance traveled decreased over time for the control group (p<0.001) while increasing over time for the CKO group (p=0.045; Figure 5C, Left graph). There was no difference in the max speed recorded between the control and CKO groups (p=0.229), but the maximum speed decreased over time for the control group (p<0.001) while there was no change for the CKO group (Figure 5D, Left graph, p=0.167).

Throughout the experiment, *Miro1^CKO^* mice displayed anxiety-like behaviors such as freezing, circling the perimeter and avoiding the center of the open field (Figure 5E,F). The control group had a smaller number of freezing episodes when compared to the CKO group (p=0.009). The number of freezing episodes seems to change over time, but it was not statistically significant (p=0.092), and there was no interaction effect between between genotype and time (p=0.598). The traces of the *Miro1^CKO^* mice consistently showed the mice circling the outside of the arena, distinct from control mice traces (Figure 5A).

To further evaluate anxiety-like behavior in the *Miro1^CKO^*mice, we employed the elevated plus maze test. Traces indicated that the *Miro1^CKO^*mice spent more time in the open arms of the elevated plus maze (Figure 6A). Interestingly, the *Miro1^CKO^*mice also showed greater variability between mice (Figure 6B). The data indicate that during the first half of the experiment both groups spent the same average amount of time in the closed and open arms (Table 1). During the second half of the experiment, the controls spent more time in the closed while the *Miro1^CKO^* mice spent more time in the open arms (Figure 6C,D). Additionally, the *Miro1^CKO^* mice had longer average visits to the open arms than controls, and the *Miro1^CKO^* mice covered less distance than the controls in the closed arms (Table 1). The overall means in the first and second half of the experiments for the controls and *Miro1^CKO^* mice were not statistically different for speed in the closed and open arms. These data were an unexpected contrast to the open field test. This led us to consider an alternative explanation, that the *Miro1^CKO^* mice manifest a different kind of anxiety related disorder, seeking to avoid closed spaces, analogous to claustrophobia.

**Figure 6:**
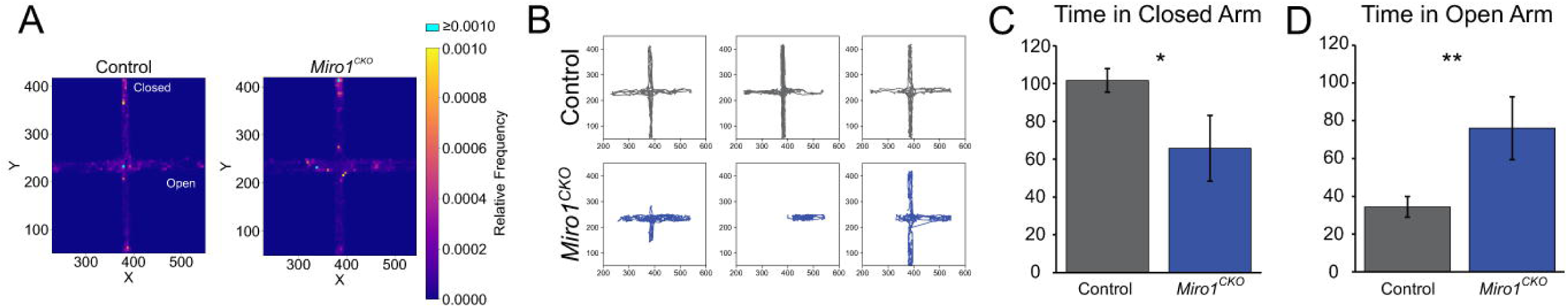
*Miro1^CKO^*mice prefer the open arm of the elevated plus maze. **A,** Coordinates from each mouse’s trace are overlayed to display where control and *Miro1^CKO^* mice spent time in the elevated plus maze. **B,** Sample traces from control and *Miro1^CKO^*mice. **C, D,** Quantification of time spent in the closed and open arm of the elevated plus maze during the second half of the experiment (150-300s; Closed Arm: t(26)=2.420, p=0.0228, Student’s Unpaired T-Test, Open Arm: t(26)=3.013, p=0.0057, Student’s Unpaired T-Test). * P≤0.05, **P≤0.01

**Table 1:** Elevated plus maze data from the first and second half of the experiment. Control and *Miro1^CKO^* data listed as mean (SEM). P-values resulted from Student’s t-tests.

|  | First Half of Experiment<br>(0-150s) |  |  | Second Half of Experiment<br>(150-300s) |  |  |
| --- | --- | --- | --- | --- | --- | --- |
| Variable | Control | *Miro1<sup>CKO</sup>* | p-value | Control | *Miro1<sup>CKO</sup>* | p-value |
| Time (s) |  |  |  |  |  |  |
| Closed Arm | 62.395 (8.580) | 67.489 (22.361) | 0.797 | 101.684 (6.206) | 65.767 (17.400) | 0.023* |
| Open Arm | 64.584 (9.735) | 67.711 (23.444) | 0.884 | 34.4 (5.493) | 76.011 (16.652) | 0.006* |
| Distance (m) |  |  |  |  |  |  |
| Closed Arm | 3.434 (0.355) | 1.943 (0.861) | 0.067 | 4.537 (0.402) | 2.077 (0.413) | <0.001* |
| Open Arm | 2.498 (0.350) | 1.307 (0.566) | 0.074 | 1.395 (0.290) | 2.212 (0.673) | 0.201 |
| Speed (m/s) |  |  |  |  |  |  |
| Closed Arm | 0.090 (0.008) | 0.071 (0.026) | 0.346 | 0.068 (0.006) | 0.069 (0.011) | 0.888 |
| Open Arm | 0.068 (0.006) | 0.049 (0.012) | 0.133 | 0.071 (0.005) | 0.055 (0.009) | 0.117 |
| Mean Visit (s) |  |  |  |  |  |  |
| Closed Arm | 19.658 (7.487) | 41.9 (22.699) | 0.233 | 30.863 (6.976) | 54.562 (21.264) | 0.182 |
| Open Arm | 14.95 (3.947) | 65.283 (28.762) | 0.012* | 7.621 (0.962) | 35.7 (19.192) | 0.021* |

### Miro1^CKO^ mice avoid confined spaces

Since the *Miro1^CKO^* mice spent more time in the open arms than the closed arms of the elevated plus maze, we hypothesized they avoided confined spaces. To further investigate this possibility, we utilized a test where mice had access to a wide space and a narrow space (Figure 7A; El-Kordi et al., 2013). *Miro1^CKO^* mice had a longer latency to enter and fewer entries into the narrow portion of the box when compared to controls (Figure 7C). Additionally, *Miro1^CKO^* mice spent significantly less time and traveled less distance in the narrow part of the box than the controls (Figure 7B,D). This data, together with the elevated plus maze data, support the hypothesis that the *Miro1^CKO^* mice avoid confined spaces.

**Figure 7:**
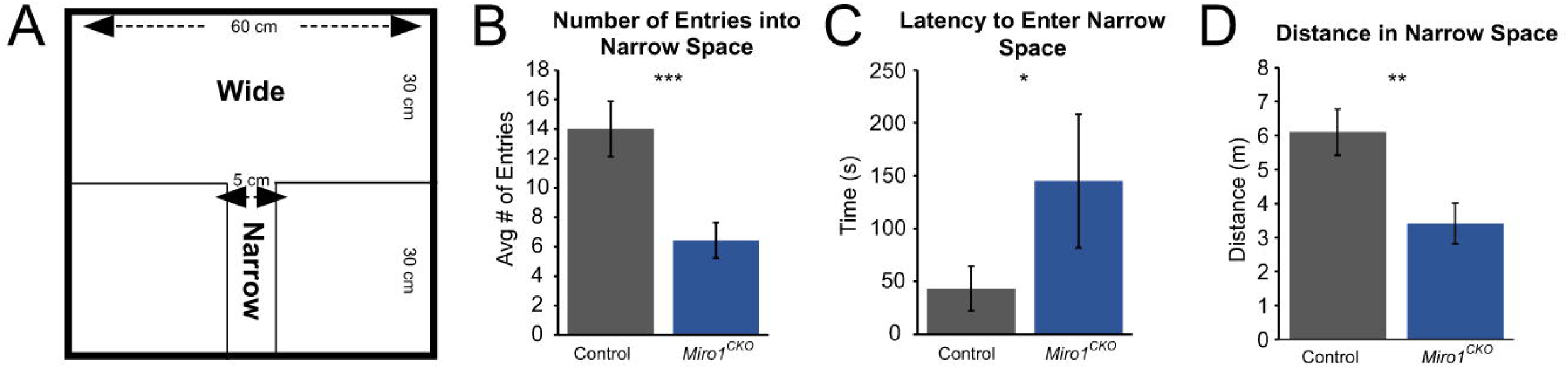
*Miro1* ablated mice avoid narrow spaces. **A,** Dimensions of the wide and narrow spaces in the box (adopted from (El-Kordi et al., 2013)). **B-D,** Quantification of the number of entries, latency to enter, and distance in the narrow space. (Controls n = 15, *Miro1^CKO^* n = 7, Latency to enter: t(20)=2.301, p=0.0323, Student’s Unpaired T-Test, Entries into narrow space: t(20)=3.935, p=0.0008, Student’s Unpaired T-Test, Distance in narrow space: t(20)=3.034, p=0.0065, Student’s Unpaired T-Test). * P≤0.05, **P≤0.01, *** P≤0.001

## Discussion

Our study reveals a unique role for *Miro1* during early cortical development and an association with an anxiety-like behavior. The abrogation of *Miro1* from excitatory neural progenitors results in a mis-localization of mitochondria to the rear of radially migrating neurons. Cycling cells labeled with KI67 are shifted with more cells at the ventricular surface at E13.5. The resulting cerebral cortex of adult *Miro1^CKO^* mice is a decreased overall brain weight and decreased cortical volume. *Miro1^CKO^*cortices show a significant decrease in the number of CTIP2 neurons in the deeper layers of the rostral forebrain as well as disorganization of CUX1-labeled neurons in layers 2/3. Furthermore, we observed selective behavioral abnormalities in the *Miro1^CKO^*mice.

In contrast to control mice, *Miro1^CKO^* mice demonstrate abnormal home cage behavior and poor self-care, including neglecting to build a nest, along with an aversion to handling. Motor testing revealed *Miro1^CKO^* mice to have mild deficits in grip strength, vertical pole, hanging wire, and wire grid; all less severe than those seen in *Eno2-cre;Miro1^CKO^* mice. Unlike the *Eno2-cre;Miro1^CKO^* mice, our *Emx1-Cre;Miro1^CKO^*mice remain able to normally move within their cage (Nguyen et al., 2014).

Open field testing found *Miro1^CKO^* mice displayed anxiety-like behaviors, spending more time on the outer edge of the arena than in the center and exhibiting increased freezing. They also have an aversion to closed spaces, as demonstrated on the elevated plus maze and the wide/narrow box test.

This is the first study to examine a role for *Miro1* in migrating excitatory neurons. Previous studies investigating a role for *Miro1* in the developing nervous system have found profound neurodegenerative symptoms including rigidity, spasticity, and death during early postnatal development, however neuronal migration was not evaluated (Nguyen et al., 2014). These studies have shown histological changes in neurite extension and degeneration, mitochondrial localization, and bunina-like body formation during development (Nguyen et al., 2014). Research from our lab suggests that mitochondrial dynamics and their ATP-producing pathways are important early in development during neuronal migration (Lin-Hendel et al., 2016). These processes appear to be neuronal subtype specific. Excitatory neurons utilize glycolysis and/or oxidative phosphorylation for energy production interchangeably during migration whereas cortical inhibitory neurons are fully dependent on oxidative phosphorylation (Lin-Hendel et al., 2016). In addition, mitochondrial dynamics differ in each neural subtype. Mitochondria in inhibitory neurons are highly dynamic through migration whereas they are primarily adjacent to the nucleus in the direction of the leading process in migrating excitatory neurons (Lin-Hendel et al., 2016). Here we find perturbing mitochondrial dynamics in migrating excitatory neurons results in subtle changes in the mature cerebral cortical architecture.

In addition to structural and molecular changes, these are the first data implicating *Miro1* in the pathogenesis of behavior phenotypes observed in neurodevelopmental disorders. While *MIRO1* has been linked to neurodegenerative disorders, its role in neurodevelopmental diseases such as autism spectrum disorders or schizophrenia has been incompletely studied (Nguyen et al., 2014; Kontou et al., 2021). Interestingly, the *Miro1^CKO^*mice display similar behaviors to Neurogranin and GPM6a knockout mice. Neurogranin has been implicated in neurodevelopmental disorders such as ADHD, autism, and schizophrenia and is a postsynaptic protein kinase that binds to calmodulin in the absence of calcium. Its expression begins in the first 3 weeks after birth and is known to be expressed in hippocampus pyramidal and granular neurons. Previous research suggests that the absence of neurogranin causes changes in synaptic plasticity including paired-pulse depression, synaptic fatigue, and long-term potentiation induction (Pak et al., 2000). Neurogranin knockout mice show decreased nesting, hyperactive behavior in their home cage, decreased time spent in the center of the open field arena, and increased time in the open arm of the elevated plus maze (Nakajima et al., 2021). Given the phenotypic similarities, studying interactions between mitochondria and neurogranin seems appropriate, or possibly neurogranin and MIRO1.

Similarly, GPM6a has been linked to autism and schizophrenia and is suggested to act as a nerve growth factor-gated calcium channel (Mukobata et al., 2002). GPM6a is known to play a role in developing neurons and is suggested to participate in neuronal migration, neuronal differentiation, and synapse development, including neurite outgrowth and spine formation (Mukobata et al., 2002; Alfonso et al., 2005; Michibata et al., 2008; Zhao et al., 2008; Mita et al., 2015; Formoso et al., 2016; Aparicio et al., 2023). Studies examining GPM6a knockout mice have found similar behavioral changes, including increased time spent in the open arm of the elevated plus maze and decreased time spent in the narrow portion of the wide/narrow box (El-Kordi et al., 2013).

Calcium is an important modulator of neurodevelopmental processes including neuronal migration and synaptic transmission. Previous research suggests that calcium regulates leading process extension and branching as well as organization during neuronal migration, acting as a “stop” and “go” signal in the developing cortex and is important for neurotransmitter release in the mature cortex (Horigane et al., 2019, review). MIRO1 has EF-1 and EF-2 calcium sensing domains that play a role in directing calcium shuttling mitochondria to regions in need of calcium buffering. Although previous research suggests *Miro1* loss does not affect the cytosolic or mitochondria calcium concentrations in mouse embryonic fibroblasts during development, ablation of *Miro1* has been found to decrease endoplasmic reticulum-mitochondrial tethering which alters calcium buffering and contributes to increasing autophagy and mitophagy leading to neuronal death in neurodegenerative diseases such as Parkinson’s and Alzheimer’s diseases (Berenguer-Escuder et al., 2020; Kam et al., 2020). One limitation of our current study is that calcium levels have not been established in developing excitatory neurons. Future studies will address the molecular and cellular underpinnings including the role of calcium dynamics in the anxiety-like behaviors that *Miro1^CKO^* mice display. Similarities to other mouse models that have indicated potential calcium signaling dysregulation make this an intriguing direction for future study.

MIRO1 is not the only mitochondrial motor adaptor protein; TRAK and MYO19 can bind to and locate mitochondrial on microtubules and actin respectively, potentially accounting for different localization patterns within a cell (MacAskill et al., 2009; López-Doménech et al., 2018; Kontou et al., 2021). It is currently not known whether these proteins are involved in certain types of localization within developing neurons and how this affects organization of, and communication between, mature neurons. Future studies are required to establish the involvement of these other mitochondrial motor adaptor proteins to further characterize their respective roles in developing neurons.

The underlying mechanisms of neuropsychiatric disorders are not well understood. This study suggests that mitochondrial location in migrating excitatory neurons could play a role in the development and onset of behavioral phenotypes in neuropsychiatric diseases such as autism spectrum disorder and schizophrenia. Our mouse model, using an *Emx1-cre* to ablate *Miro1,* displays distinct anxiety-like behavior patterns that are distinct from those observed in previous models such as when *Miro1* is removed using the *Eno2-cre.* The latter model showed classic characteristics of neurodegeneration including muscle spasticity and weakness, hindlimb clasping, premature death around postnatal day 40, and pathologic features such as bunina bodies (Nguyen et al., 2014). Unfortunately, their analysis of P30 cerebral cortex was limited and developmental changes were not considered. One possible explanation for the difference between our model and the *Eno2-Cre;Miro1* mutant is that each disrupts specific networks of neurons that leads to different disease phenotypes; some that have an early onset and others that have a later onset. Distinguishing the networks of neurons that are impacted by the ablation of *Miro1* in each of these cases could provide a deeper understanding of the underlying circuitry dysfunction that plagues individuals with neurodevelopmental and neurodegenerative diseases. This may also lead to novel therapies that could improve the outcomes and quality of life for afflicted individuals. Finally, our study provides a new model to investigate the neurobiology underlying behavioral phenotypes related to anxiety-like disorders.

Evidence of mitochondrial dysfunction has been recognized in neuropsychiatric conditions such as autism spectrum disorder and schizophrenia (Poling et al., 2006; DiMauro and Schon, 2008; Weissman et al., 2008). Our study suggests that early loss of *Miro1* in migrating excitatory neurons could contribute to and sustain behavioral symptoms that accompany these disorders. A continued focus and understanding of the cellular and molecular mechanisms of *Miro1* could lead to better targeting of treatments and a deeper knowledge of the origins of neuropsychiatric disorders.

## Author contributions

A.K.M., G.C. and J.A.G. designed research; A.K.M., M.S., C.R., C.F., L.J., R.S., M.C., C.C., L.B., S.S., N.K., J.G, G.C. performed research; A.K.M., M.S., C.R., C.F, L.J., L.B., S.S., S.K., and M.T. analyzed data; A.K.M. wrote the paper. The authors declare no competing financial interests.

## Acknowledgements

This work was supported by the National Institutes of Neurologic Disorders and Stroke (NS100007 to JAG) and financial support from the Hamilton College Biology and Psychology Departments and Dean’s Office. We would like to thank Tom Freeland and Walt Zarnoch for giving their time to build our elevated plus maze and the wide-narrow box, Trevor Lancon for helping to compute the mitochondria localization data, Siobhan Robinson for the use of her microscope and microtome, the Masonic Medical Research Institute especially Chase Kessinger for the use of their cryostat, and Russell Hardesty for his help with constructing traces, heat maps, and graphs. We would also like to thank members of the Golden Lab for many discussions, suggestions, and support. We would specifically like to thank Brenna Stallings for managing the mouse colony at Brigham and Women’s Hospital and providing genotyping results for each of the litters.

